# Bayesian weighing of electron cryo-microscopy data for integrative structural modeling

**DOI:** 10.1101/113951

**Authors:** Massimiliano Bonomi, Samuel Hanot, Charles H. Greenberg, Andrej Sali, Michael Nilges, Michele Vendruscolo, Riccardo Pellarin

## Abstract

**Summary:** Cryo-electron microscopy (cryo-EM) has become a mainstream technique for determining the structures of complex biological systems. However, accurate integrative structural modeling has been hampered by the challenges in objectively weighing cryo-EM data against other sources of information due to the presence of random and systematic errors, as well as correlations, in the data. To address these challenges, we introduce a Bayesian scoring function that efficiently and accurately ranks alternative structural models of a macromolecular system based on their consistency with a cryo-EM density map and other experimental and prior information. The accuracy of this approach is benchmarked using complexes of known structure and illustrated in three applications: the structural determination of the GroEL/GroES, RNA polymerase II, and exosome complexes. The approach is implemented in the open-source *Integrative Modeling Platform* (http://integrativemodeling.org), thus enabling integrative structure determination by combining cryo-EM data with other sources of information.

**Highlights:**

- We present a modeling approach to integrate cryo-EM data with other sources of information
- We benchmark our approach using synthetic data on 21 complexes of known structure
- We apply our approach to the GroEL/GroES, RNA polymerase II, and exosome complexes

## Introduction

Over the last two decades, cryo-electron microscopy (cryo-EM) has enabled the structural characterization of complex biological systems beyond the capabilities of traditional techniques, such as X-crystallography and nuclear magnetic resonance (NMR) spectroscopy (Callaway, 2015; Kuhlbrandt, 2014; Nogales, 2016). This progress has been fueled by the continuous advances in both instrumentation and software for cryo-EM image processing (Bai et al., 2015; Glaeser, 2016; Li et al., 2013). As a result, the resolution of the structures from cryo-EM is rapidly approaching that of X-ray crystallography. Most importantly, cryo-EM does not require crystallizing the system prior to data acquisition, needs a small amount of sample, does not require isotopic labeling, and is applicable to systems larger than ~100 kDa. Furthermore, cryo-EM has the potential of identifying multiple different structural states in a single experiment (Bai et al., 2015; Callaway, 2015; Glaeser, 2016; Nogales, 2016), provided that they can be disentangled during image classification.

A number of approaches have been proposed to model macromolecular structures based on cryo-EM density maps (Lopez-Blanco and Chacon, 2015; Schroder, 2015). Generally speaking, these techniques can use one or more of the following strategies: rigid-body fitting of components of known structures, flexible refinement, use of homology modeling or *de novo* protein structure prediction of the components, and integrative modeling based on multiple types of experimental data. The most popular software packages for cryo-EM-based modeling include Chimera (Pettersen et al., 2004), EMfit (Rossmann et al., 2001), Modeller (Sali and Blundell, 1993), SITUS (Wriggers, 2012), MultiFit (Lasker et al., 2009), EMFF (Zheng, 2011), MDFF (Trabuco et al., 2008), Flex-EM (Topf et al., 2008), γ-TEMPy (Pandurangan et al., 2015), COAN (Volkmann and Hanein, 1999), MDFIT (Ratje et al., 2010), Fold-EM (Saha and Morais, 2012), ROSETTA (DiMaio et al., 2009), EM-fold (Lindert et al., 2012), IMP (Russel et al., 2012), RELION (Scheres, 2012a), ISD (Habeck, 2017), and Phenix (Adams et al., 2011). The majority of these approaches generate structural models that minimize the deviation between observed and predicted cryo-EM density maps, including by molecular dynamics (MD), Monte Carlo (MC) or normal modes analysis techniques (Lopez-Blanco and Chacon, 2015).

Several methods have been developed with the purpose of fitting the components of large macromolecular complexes into low-resolution density maps. A subset of these of methods use scoring functions based on cross-correlation (CC) or Laplacian-filtered CC between a target map and a simulated map, sampling using three-dimensional (3D) Cartesian FFT coupled with exhaustive rotational samples, such as COLORES (Chacon and Wriggers, 2002), gEMfitter (Hoang et al., 2013), and PowerFit (van Zundert and Bonvin, 2015). These methods are normally used incrementally, i.e. by fitting one subunit at the time. In contrast, other modeling software packages simultaneously assemble multiple components of the complex.ATTRACT-EM (de Vries and Zacharias, 2012), for example, uses Gaussians positioned at the center of each voxel of the map, a coarse-grained representation of the model structure, and a gradient vector matching as energy function.

Despite the success of these methods, the translation of cryo-EM density maps into structural models still presents several challenges, especially in integrative structural modeling, where cryo-EM data are combined with other sources of information. First, cryo-EM density maps are affected by random and systematic errors (Bonomi et al., 2017; Schneidman-Duhovny et al., 2014). In particular, radiation damage to the sample upon prolonged exposure to the electron beam often results in regions of the density map at resolution lower than the average. Second, despite progress in methods for 2D classification and 3D reconstruction, the final maps might still average out images of particles in different conformations (Bonomi et al., 2016). Finally, cryo-EM maps are typically defined by a set of data points, or voxels, representing the electron density on a grid in real space. Neighbouring voxels do not provide independent information on the system, but instead are affected by a certain degree of spatial correlation. Accounting for correlation as well as the presence of noise in the data is crucial when integrating cryo-EM with other experimental data (Ward et al., 2013), as the information and noise content of each piece of data needs to be accurately quantified to avoid biasing a model (Schneidman-Duhovny et al., 2014).

Here, we introduce a Bayesian approach (Rieping et al., 2005) to model the structure of a macromolecular system by optimally combining cryo-EM data with other input information. Bayesian inference and maximum-likelihood methods are not novel to the cryo-EM field (Scheres, 2012b; Sigworth et al., 2010), as they were initially introduced for aligning structurally homogenous sets of 2D images (Sigworth, 1998), and they are now widely used by software packages such as RELION (Scheres, 2012a) for single-particle reconstruction. In our approach, we use Bayesian inference in a way similar to that discussed in a recent paper (Habeck, 2017), i.e. to determine the optimal weight of cryo-EM data in integrative structural modeling.

Our approach models the structure of the system while simultaneously and automatically quantifying the level of noise in the data. Furthermore, the input data are represented in terms of a Gaussian mixture model (GMM) (de Vries and Zacharias, 2012; Jonic et al., 2016; Kawabata, 2008; Robinson et al., 2015), rather than using the standard voxel representation. This procedure has several advantages: a) it alleviates the problem of voxel correlation by decomposing the density map into a set of nearly-independent GMM components; b) it is computationally efficient; and c) it enables a multi-scale representation of the model, from coarse-grained for initial efficient sampling to atomistic for refinement of high-resolution maps. By accounting for both data noise and correlation, this approach enables an effective use of cryo-EM density maps in integrative structural modeling.

In the following, we first outline our modeling approach and then benchmark its accuracy using synthetic low-resolution data of several protein/DNA complexes. Finally, we apply our approach to the integrative modeling of the GroEL/ES complex, as well as the RNA polymerase II and the exosome complexes, in which we combine cryo-EM with chemical cross-linking/mass spectrometry (XL-MS) data. This method is implemented in the open-source Integrative Modeling Platform (IMP) package (http://integrativemodeling.org) (Russel et al., 2012), thus enabling integrative structure determination of biological systems based on a variety of experimental data, including FRET and NMR spectroscopies, XL-MS, small angle X-ray scattering, and various proteomics data.

## Results

### Protocol for low-resolution modeling of cryo-EM density maps

We implemented in IMP (Russel et al., 2012) a pipeline that enables the multi-scale modeling of macromolecular structures based on cryo-EM data and other structural information, given partial knowledge of subunits structures. The details of our approach are illustrated in the STAR Methods. The general 4-stage protocol proceeds as follows (**Fig.1**):

1. Gather the data, including the sequences of subunits, their structures (*eg*, from X-ray crystallography, NMR spectroscopy, homology modeling, and *ab initio* prediction), and the target cryo-EM density map (**Fig.1.1**).
2. Convert the data into a scoring function for ranking alternative structural models:

A. Generate a GMM representation of the density map (data-GMM) by using a divide-and-conquer algorithm (**Fig.1.2A, Fig.2, and Fig.S1**).
B. Assign a representation to the different components of the complex (**Fig.1.2B**). Subunits are represented by spherical beads to coarse-grain the atomic degrees of freedom. For a given domain, the beads are either constrained into a rigid body or allowed to move flexibly, depending on the uncertainty about the domain structure. The beads represent one or more contiguous residues, depending on the level of coarse-graining (Erzberger et al., 2014; Fernandez-Martinez et al., 2016; Robinson et al., 2015).
C. The electron density of the model is also described by a GMM (model-GMM) and is used to compute the fit of the model to the cryo-EM density map (**Fig.1.2C**).
D. The scoring function that ranks the models according to how well they fit the input information is derived from the posterior probability, which includes a likelihood function for the cryo-EM data (**Fig.S2**), and prior terms such as the bead sequence connectivity and excluded volume (**Fig.1.2D**).
3. Sample models using MC and replica exchange methods (Swendsen and Wang, 1986), with an iterative approach to maximize sampling exhaustiveness (**Fig.1.3 and Fig.S3**).
4. Analyze the sampled models in terms of their variability by clustering. (**Fig.1.4**).

**Figure 1.**
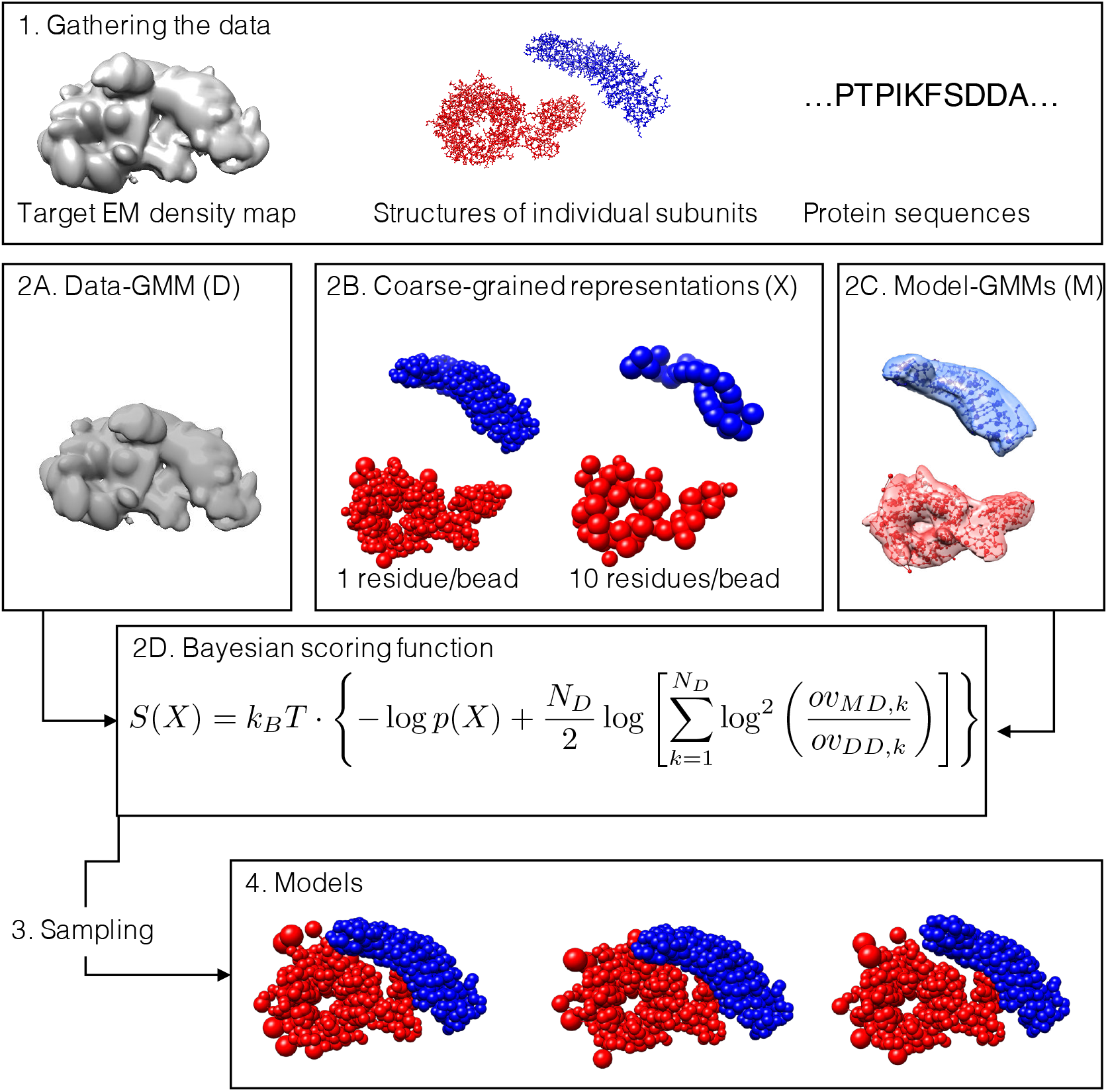
Workflow for multi-scale modeling of cryo-EM data. (1) The input information for the modeling protocol consists of: an experimental cryo-EM density map (left), the structures of the subunits (center), and the sequences of the subunits (right). (2A) The density map is fitted with a GMM *(ie*, the data-GMM) using our divide-and-conquer approach. (2B) The atomistic coordinates of the subunits are suitably coarse-grained into large beads. Regions without a known atomistic structure are represented by a string of large beads, each representing a set of residues. (2C) GMM for the subunits (ie, the model-GMMs) are also computed from the atomistic coordinates. (2D) The Bayesian scoring function encodes prior information about the system and measures the agreement between the data-GMM and the model-GMM. (3) Structural models are sampled by MC coupled with replica exchange, with or without the iterative sampling protocol. (4) The generated models are analysed.

**Figure 2.**
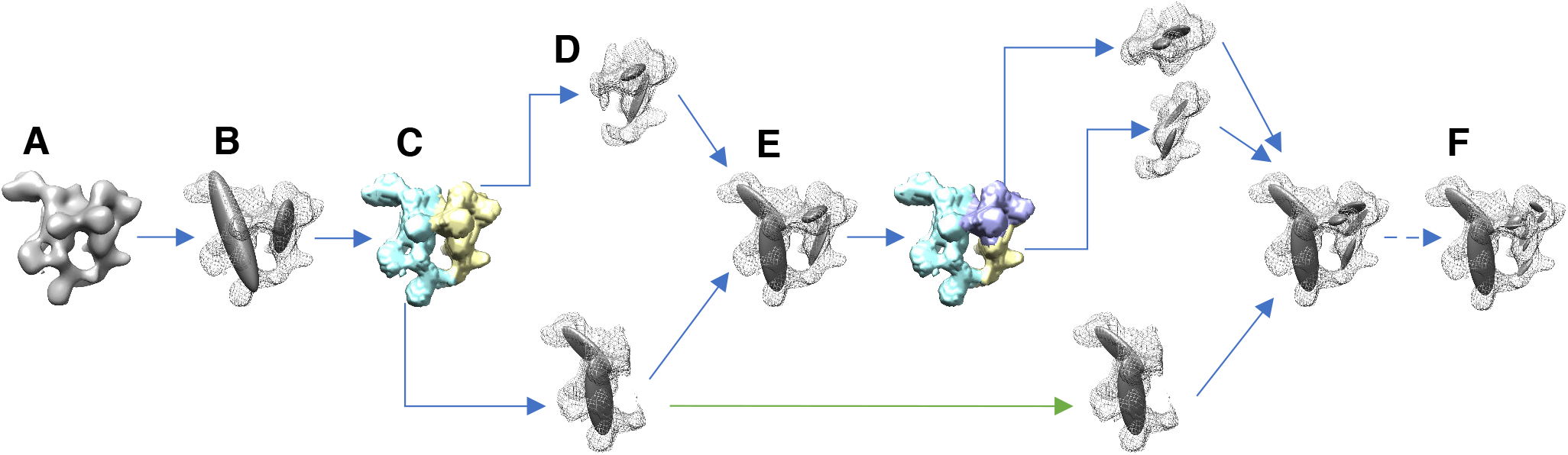
Divide-and-conquer approach for fitting cryo-EM density maps with a Gaussian mixture model (GMM) (A) The input map is thresholded according to the recommended threshold. (B) The resulting map is initially fitted using a GMM with 2 components. (C) Each component of the GMM is used to partition the map into overlapping sub-maps. (D) Each sub-map is fitted using a GMM with 2 components, similarly to step B. (E) The sum of all the GMMs of the sub-maps results in a data-GMM that approximates the original map. The accuracy of approximation increases at every iteration. (F) The fitting procedure is iterated until the data-GMM reaches an optimal accuracy. The green arrow indicates a branch that was stopped because the local CC was higher than 0.95.

**Figure 3.**
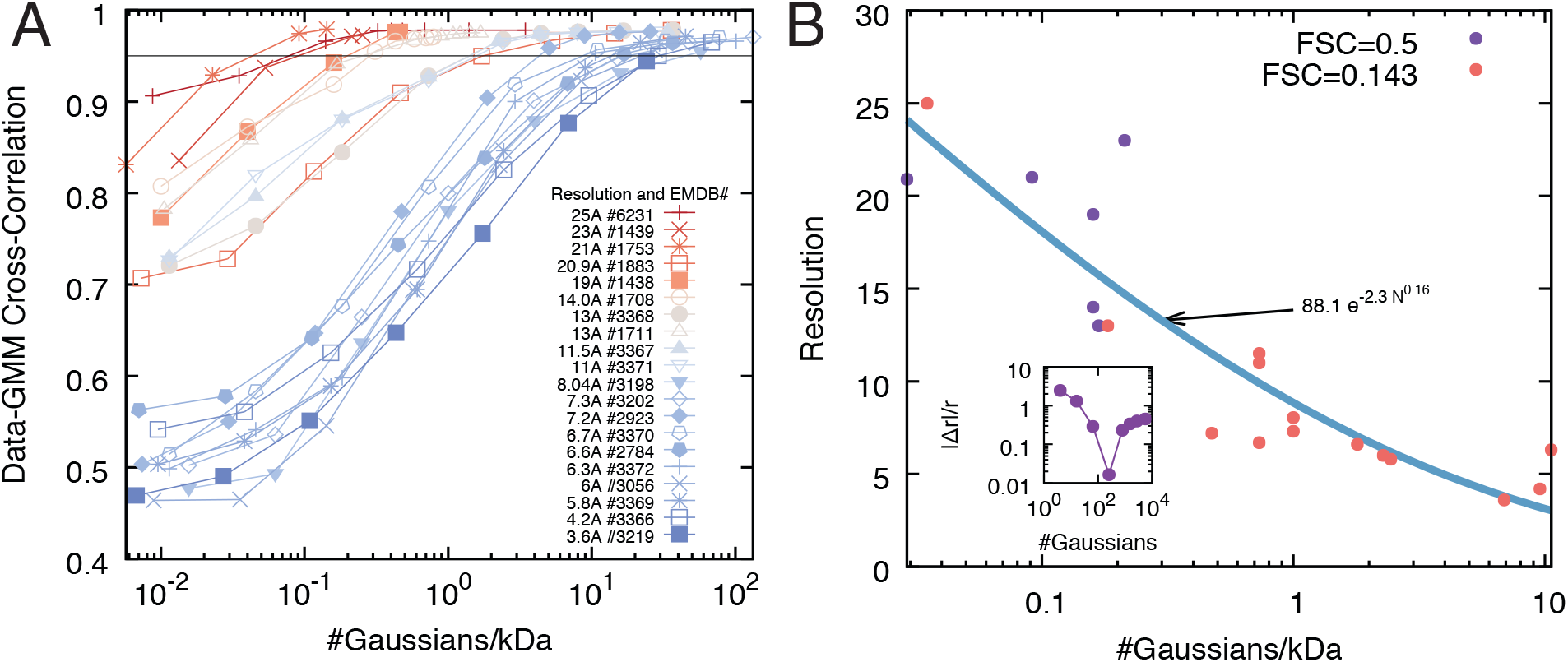
Benchmark of the divide-and-conquer fit of the data-GMM. (A) The accuracy of the divide-and-conquer approach is measured using the correlation coefficient between the input map and the corresponding data-GMMs obtained at different iterations. The accuracy increases with the number of components in the mixture, and the saturation point (*ie*, the number of components beyond which the accuracy does not increase significantly) depends on the resolution of the experimental map (red and blue curves are low and high resolutions, respectively). (B) Relationship between map resolution and number of components of the data-GMM. For all the density maps of panel A, the experimental resolution is plotted as a function of the optimal number of components of the data-GMM normalized by the molecular weight of the complex (solid circles). The points are fitted using a power law (blue line). The orange and purple circles correspond to maps whose resolution was determined by the Fourier Shell Correlation 0.143 and 0.5, respectively. (B, inset) For each density map, the optimal number of components is computed as the minimal absolute relative deviation *| Δr| /r* between the data-GMM resolution and the density map resolution.

**Figure 4.**
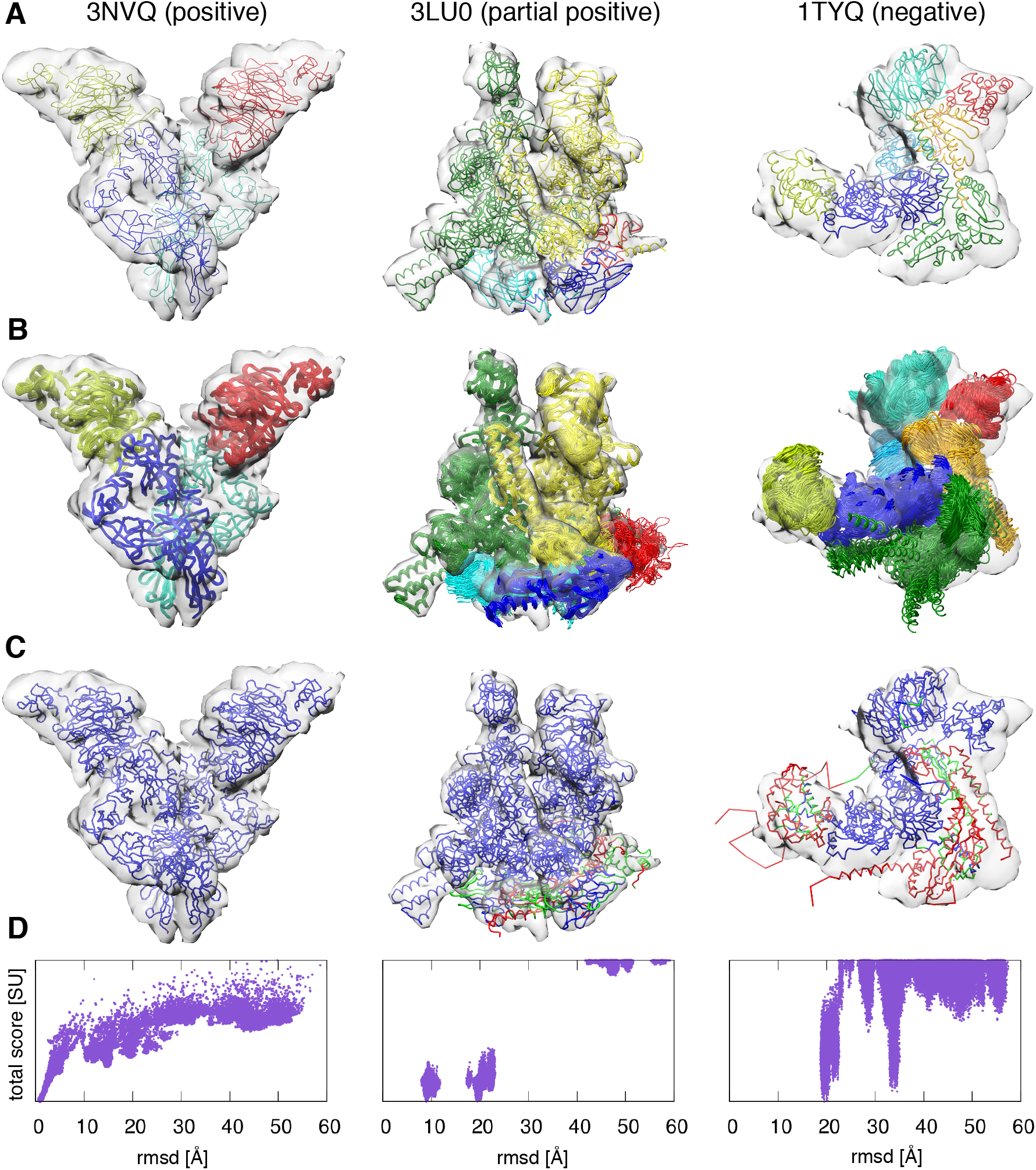
Benchmark of the modeling protocol. Examples of each of the three possible outcomes of the benchmark: positive (first column, PDB code 3NVQ), partial positive (second column, PDB code 3LU0), and negative (third column, PDB code 1TYQ). (A) Native structures and simulated 10 Å resolution cryo-EM density maps. (B) 50 best scoring models displayed with the simulated cryo-EM density maps. (C) Residue-wise accuracy of the best scoring models: residues whose positions deviate from the native structure less than 10 Å, between 10 and 20 Å, and above 20 Å are coloured in blue, green, and red, respectively. (D) Total score of all the sampled models as a function of the total rmsd from the native structure.

### Benchmark of the divide-and-conquer fit of the data-GMM

We assessed the accuracy of our divide-and-conquer approach by determining the data-GMM of 20 experimental density maps at different resolutions (Liu et al., 2016; Malet et al., 2010; Wang et al., 2007), ranging from 3.6 Å to 25 Å (**Table 1**). This benchmark revealed that the number of Gaussian components needed to achieve a given accuracy of the optimal data-GMM varies with the resolution of the map and molecular weight of the complex (**Fig.3A**). Indeed, for a given number of components and molecular weight, the data-GMM correlation coefficient is lower for higher-resolution maps. In other words, high resolution maps and maps of high molecular weight complexes contain more information and therefore require additional components to describe the ensemble of their features.

**Table 1.** Benchmark of the divide-and-conquer approach for GMM fitting. For each of the system studied, we report: the EMDB accession code, the reference paper, the molecular weight of the complex, the resolution of the map, the method used to quantify the experimental resolution, the optimal number of components, the resolution of the optimal GMM, the CC between the experimental cryo-EM map and the optimal GMM.

| EMDB<br>code | reference | molecular<br>weight<br>[kDa] | resolution<br>[Å] | resolution<br>method | optimal<br>number of<br>components | GMM<br>resolution<br>[Å] | CC |
| --- | --- | --- | --- | --- | --- | --- | --- |
| 1439 | (Wang et al., 2007) | 300 | 23.0 | FSC 0.5 | 64 | 21.6 | 0.97 |
| 1438 | (Wang et al., 2007) | 400 | 19.0 | FSC 0.5 | 184 | 19.5 | 0.97 |
| 1708 | (Malet et al., 2010) | 400 | 14.0 | FSC 0.5 | 304 | 14.7 | 0.97 |
| 3368 | (Liu et al., 2016) | 350 | 13.0 | FSC 0.143 | 256 | 10.8 | 0.93 |
| 3367 | (Liu et al., 2016) | 350 | 11.5 | FSC 0.143 | 784 | 11.6 | 0.97 |
| 3371 | (Liu et al., 2016) | 350 | 11.0 | FSC 0.143 | 844 | 10.5 | 0.96 |
| 3370 | (Liu et al., 2016) | 350 | 6.7 | FSC 0.143 | 1023 | 6.5 | 0.92 |
| 3372 | (Liu et al., 2016) | 350 | 6.3 | FSC 0.143 | 1023 | 5.7 | 0.90 |
| 3369 | (Liu et al., 2016) | 420 | 5.8 | FSC 0.143 | 1024 | 6.2 | 0.85 |
| 3366 | (Liu et al., 2016) | 420 | 4.2 | FSC 0.143 | 12219 | 3.9 | 0.95 |
| 6231 | (Booth et al., 2014) | 457 | 25.0 | FSC 0.143 | 64 | 24.2 | 0.97 |
| 1753 | (Vannini et al., 2010) | 700 | 21.0 | FSC 0.5 | 100 | 24.0 | 0.98 |
| 1883 | (Czeko et al., 2011) | 550 | 20.9 | FSC 0.5 | 64 | 19.1 | 0.82 |
| 1711 | (Julian et al., 2011) | 380 | 13.0 | FSC 0.5 | 640 | 13.9 | 0.98 |
| 3198 | (Fernandez-Leiro et al., 2015) | 256 | 8.0 | FSC 0.143 | 256 | 7.8 | 0.78 |
| 2923 | (Martinez-Rucobo et al., 2015) | 540 | 7.2 | FSC 0.143 | 1012 | 6.8 | 0.90 |
| 2784 | (Plaschka et al., 2015) | 570 | 6.6 | FSC 0.143 | 1024 | 6.9 | 0.84 |
| 3056 | (des Georges et al., 2015) | 450 | 6.0 | FSC 0.143 | 1024 | 6.2 | 0.83 |
| 3202 | (Fernandez-Leiro et al., 2015) | 256 | 7.3 | FSC 0.143 | 256 | 7.5 | 0.80 |
| 3219 | (Bernecky et al., 2016) | 590 | 3.6 | FSC 0.143 | 14225 | 3.6 | 0.94 |

We used this benchmark to calculate the resolution of the cryo-EM density maps as a function of the number of Gaussians per mass-unit of the optimal data-GMM (**Fig.3B**). This relationship can be used to: a) estimate the resolution of a GMM generated from a known structure, and b) estimate the number of Gaussians needed to fit a cryo-EM density map of a given mass and resolution.

Our divide-and-conquer approach allowed us to overcome the computational inefficiency of the traditional expectation-maximization algorithm for fitting GMM with a large number of components. For example, in the case of the yeast cytoplasmic exosome at 4.2 Å resolution (EMDB code 3366) (Liu et al., 2016), our approach required 24 minutes and less than 1 GB to generate GMMs with 4, 16, 64, 256, 1024, and 4096 components. In contrast, a serial implementation on a single computer required over 48 hours and 182 GB of memory for fitting with 4096 components.

### Benchmark of the modeling protocol

We assessed the accuracy of the modeling protocol using a benchmark of 21 protein/DNA complexes consisting of 2 to 7 subunits (**Table 2**) (Velazquez-Muriel et al., 2012) and simulated cryo-EM density maps with a resolution of ~10 Å. No additional experimental data beside the crystal structure of the individual components were included, as our aim was to explore the performance of the cryo-EM scoring function alone. The detailed results of the benchmark are reported in **Table S1**.

**Table 2.** Results of the benchmark of the modeling protocol. For each of the system studied, we report: the PDB accession code, the reference paper, the number of subunits, the number of clusters, the average time needed to produce one model, the rank and rmsd of the model with minimum rmsd with respect to the reference structure (best rmsd model). We also report the following information about the best scoring model: rmsd, p(10), the data-model correlation coefficient (CC), and the average placement scores (APS) with respect to the reference structure.

| PDB<br>code | reference | #<br>subuni<br>ts | #<br>clus<br>ters | time /<br>frame<br>[s] | best rmsd<br>model |  | best scoring model |  |  |  |  |
| --- | --- | --- | --- | --- | --- | --- | --- | --- | --- | --- | --- |
|  |  |  |  |  | rank | rmsd<br>[Å] | rmsd<br>[Å] | p(10) | CC | APS [Å,°] |  |
| 2UZX | (Niemann et al., 2007) | 2 | 1 | 1.6 | 551 | 1.1 | 1.5 | 1.00 | 0.90 | 0.7 | 2.9 |
| 3R5D | (Schnell et al., 2012) | 3 | 1 | 1.1 | 63 | 1.4 | 1.9 | 1.00 | 0.93 | 1.1 | 2.6 |
| 1CS4 | (Tesmer et al., 2000) | 3 | 1 | 0.7 | 159 | 2.6 | 3.5 | 1.00 | 0.79 | 0.8 | 1.1 |
| 2WVY | (Zhu et al., 2010) | 3 | 1 | 3.0 | 480 | 0.9 | 1.3 | 1.00 | 0.98 | 0.5 | 0.6 |
| 2DQJ | (Shiroishi et al., 2007) | 3 | 3 | 0.4 | 89 | 2.0 | 2.5 | 1.00 | 0.93 | 1.6 | 3.0 |
| 1VCB | (Stebbins et al., 1999) | 3 | 1 | 0.2 | 453 | 1.8 | 2.2 | 1.00 | 0.82 | 1.0 | 2.7 |
| 2GC7 | unpublished | 4 | 1 | 0.6 | 817 | 1.3 | 2.0 | 1.00 | 0.94 | 0.9 | 1.8 |
| 2BO9 | (Pallares et al., 2005) | 4 | 1 | 0.8 | 502 | 1.3 | 1.6 | 1.00 | 0.95 | 0.9 | 1.4 |
| 2BBK | (Chen et al., 1998) | 4 | 1 | 0.9 | 857 | 2.1 | 2.4 | 1.00 | 0.90 | 1.7 | 1.0 |
| 1GPQ | (Abergel et al., 2007) | 4 | 1 | 0.5 | 805 | 1.7 | 2.2 | 1.00 | 0.90 | 1.4 | 1.5 |
| 3V6D | (Das et al., 2012) | 4 | 2 | 1.0 | 686 | 1.6 | 2.1 | 1.00 | 0.92 | 1.0 | 5.2 |
| 3SFD | (Zhou et al., 2011) | 4 | 1 | 1.3 | 385 | 1.4 | 1.5 | 1.00 | 0.94 | 0.7 | 1.0 |
| 3PDU | (Zhang et al., 2014) | 4 | 1 | 1.3 | 802 | 1.3 | 1.6 | 1.00 | 0.93 | 1.3 | 0.9 |
| 3NVQ | (Liu et al., 2010) | 4 | 1 | 2.1 | 903 | 0.9 | 1.0 | 1.00 | 0.97 | 0.6 | 0.7 |
| 2Y7H | (Kennaway et al., 2009) | 5 | 1 | 1.9 | 465 | 1.7 | 2.2 | 1.00 | 0.88 | 1.9 | 1.5 |
| 1SUV | (Cheng et al., 2004) | 6 | 1 | 1.4 | 247 | 5.2 | 5.3 | 1.00 | 0.77 | 5.2 | 0.4 |
| 1Z5S | (Reverter and Lima, 2005) | 4 | 1 | 0.3 | 892 | 8.7 | 9.0 | 0.86 | 0.87 | 1.8 | 32.1 |
| 3LU0 | (Opalka et al., 2010) | 5 | 2 | 5.1 | 800 | 9.0 | 9.3 | 0.85 | 0.74 | 4.3 | 5.7 |
| 1MDA | (Chen et al., 1992) | 6 | 2 | 1.8 | 4286 | 7.8 | 8.1 | 0.85 | 0.75 | 3.2 | 34.0 |
| 3PUV | (Oldham and Chen, 2011) | 5 | 1 | 3.1 | 698 | 22.7 | 23.4 | 0.64 | 0.74 | 8.0 | 31.1 |
| 1TYQ | (Nolen et al., 2004) | 7 | 3 | 5.1 | 599 | 19.1 | 19.8 | 0.65 | 0.65 | 4.1 | 58.2 |

The average accuracy p(10) (STAR Methods) of the whole benchmark was 88%. We classified the outcomes of our benchmark into three categories. We defined a *full positive* result when the global root mean square deviation (rmsd) with respect to the reference structure along with the rmsds of all the individual subunits were lower than 10 Å. A *partial positive* result was achieved when the global rmsd was lower than 10 Å but some of the subunits were misplaced, resulting in a rmsd greater than 10 Å for at least one subunit. A *negative* result was obtained when the global rmsd was greater than 10 Å. Out of the 21 complexes, we obtained 16 full positives (2UZX, 3R5D, 1CS4, 2WVY, 2DQJ, 1VCB, 2GC7, 2BO9, 2BBK, 1GPQ, 3V6D, 3SFD, 3PDU, 3NVQ, 2Y7H, and 1SUV), 3 partial positives (1Z5S, 3LU0, and 1MDA), and 2 negatives (3PUV and 1TYQ) (**Table 2 and Fig.4**).

The majority of the complexes belonged to the full positive category and were accurately modeled, with an average global rmsd from the reference structure equal to 2.2 Å. In the following, we discuss the few partial positive and negative results, highlighting the reasons behind their lower accuracy.

The best-scoring model of the 4-subunits 1Z5S had a rmsd of 9.0 Å, p(10) of 0.86, APS of (1.8 Å, 32.1°), and CC of 0.87. All best scoring models are grouped into a single cluster. The origin of the inaccuracy was subunit B, which was mis-rotated by almost 180°. The reason was that this subunit has a cylindrical shape and therefore the expected density is nearly invariant under rotations around the main axis.

The best-scoring model of the 5-subunits 3LU0 had a rmsd of 9.3 Å, p(10) of 0.85, APS of (4.3 Å, 5.7°), and CC of 0.74. The best scoring models were grouped into two clusters. In the first cluster containing the best scoring model, the lower accuracy of the models was due to subunits A, B, and E. Subunits A and B, while positioned in the correct region of the density map, were displaced by 10.9 Å and 7.3 Å and mis-rotated by 33.7° and 24.1°, respectively. Subunit E was also displaced by 13.5 Å and mis-rotated by 34.6°. In the second cluster, the situation was similar, with subunit E displaced even farther apart in the incorrect region of the density, with a Placement Score of (72.6 Å, 151.7°).

The best-scoring model of the 6-subunits 1MDA had a rmsd of 8.1 Å, p(10) of 0.85, APS of (3.2 Å, 34.0°), and CC of 0.75. All best scoring models were grouped into a single cluster. The origin of the inaccuracy was the orientations of subunit A and M, which were both mis-rotated by almost 180°. The reason was that these subunits have near cylindrical shape and therefore their expected densities are almost invariant under rotations around the main axis.

The best-scoring model of the 5-subunits 3PUV had a rmsd of 23.4 Å, p(10) of 0.64, APS of (8.0 Å, 31.1°), and CC of 0.74. All best scoring models were grouped into a single cluster. In the best-scoring model, subunits E, F, and G were correctly positioned. Subunits A and B were instead both misplaced and mis-rotated, with APS of (17.6 Å, 17.7°) and (18.1 Å, 136.7°), respectively. The reason was that these subunits formed a closed dimer of roughly cubical shape, whose density could be fit also by an incorrect model in which the positions of the domain of subunits A and B were swapped.

The best-scoring model of the 7-subunits 1TYQ had a rmsd of 19.8 Å, p(10) of 0.65, APS of (4.1 Å, 58.2°), and CC of 0.65. The best scoring models were grouped into 3 clusters. In all clusters, subunits D and F were both misplaced and mis-rotated. The most likely reason for this inaccuracy was that these subunits formed an elongated helical bundle of about 40 residues in length, making sampling more challenging due to steric effects. In the first cluster containing the best scoring model, subunit E was also mis-rotated by almost 180°, due to its globular shape.

### Modeling of the GroEL/ES complex

The ADP-bound GroEL/ES is a 21-subunit molecular chaperone that assists protein folding in bacteria. We used cryo-EM data at 23.5 Å resolution (EMDB code 1046) (Ranson et al., 2001) and the crystallographic structures of the subunits (PDB code 1AON) (Xu et al., 1997) (**Fig.S4**).

The 100 best-scoring models grouped into 3 clusters, which were mainly different in the orientation of the GroEL-trans subunit (**Fig.5 and Table S2**). All three clusters presented a misrotation of the GroES subunit, which was due to the small size of the subunit and the low resolution of the map (de Vries and Zacharias, 2012; Habeck, 2017; Kawabata, 2008). The rmsd of the best scoring model (**Fig.S4**) with respect to the reference structure was 9.0 Å, with p(10) of 0.98 and data-model CC of 0.85. Notably, GroES and GroEL-cis proteins were determined with lower precision than GroEL-trans.

**Figure 5.**
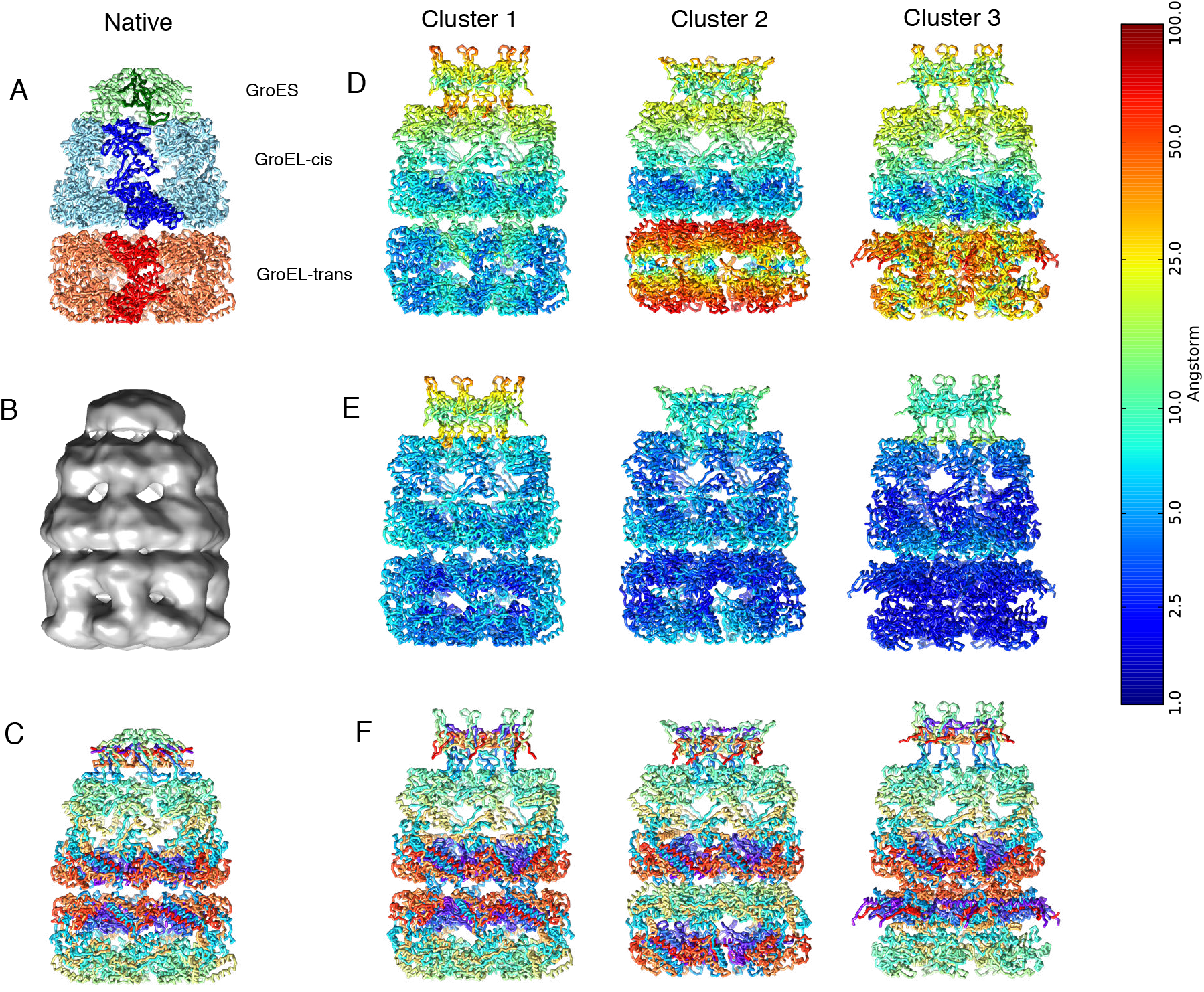
Modeling of the GroEL/ES complex. (A) Native structure of the GroEL/ES complex (PDB code 1AON). (B) Cryo-EM density map of GroEL/ES (EMDB 1046). (C) Residue indexes are color-coded using a rainbow palette, where the N-terminus is violet, the C-terminus is red, and intermediate residues are green and yellow. The three columns on the right are the representative structures of the three best-scoring clusters color coded using the rmsd from the native structure per residue (D), the per-residue precision (E), and the same color coding as in (C) to emphasize the orientation of the subunits. The color bar on the left refers to the panels (D) and (E).

### Integrative modeling of the RNA polymerase II

The yeast RNA polymerase II is a 12-subunit complex that catalyzes DNA transcription to synthesize mRNA strands (Armache et al., 2005). To model this complex, we used the structures of all its subunits as determined in the RNA polymerase II X-ray structure (PDB code 1WCM) (Armache et al., 2005). We incorporated a low-resolution cryo-EM map of the RNA polymerase II-Iwr1 complex (EMDB code 1883) (Czeko et al., 2011) and two XL-MS datasets (**Fig.6, Fig.S5, and Fig.S6**).

**Figure 6.**
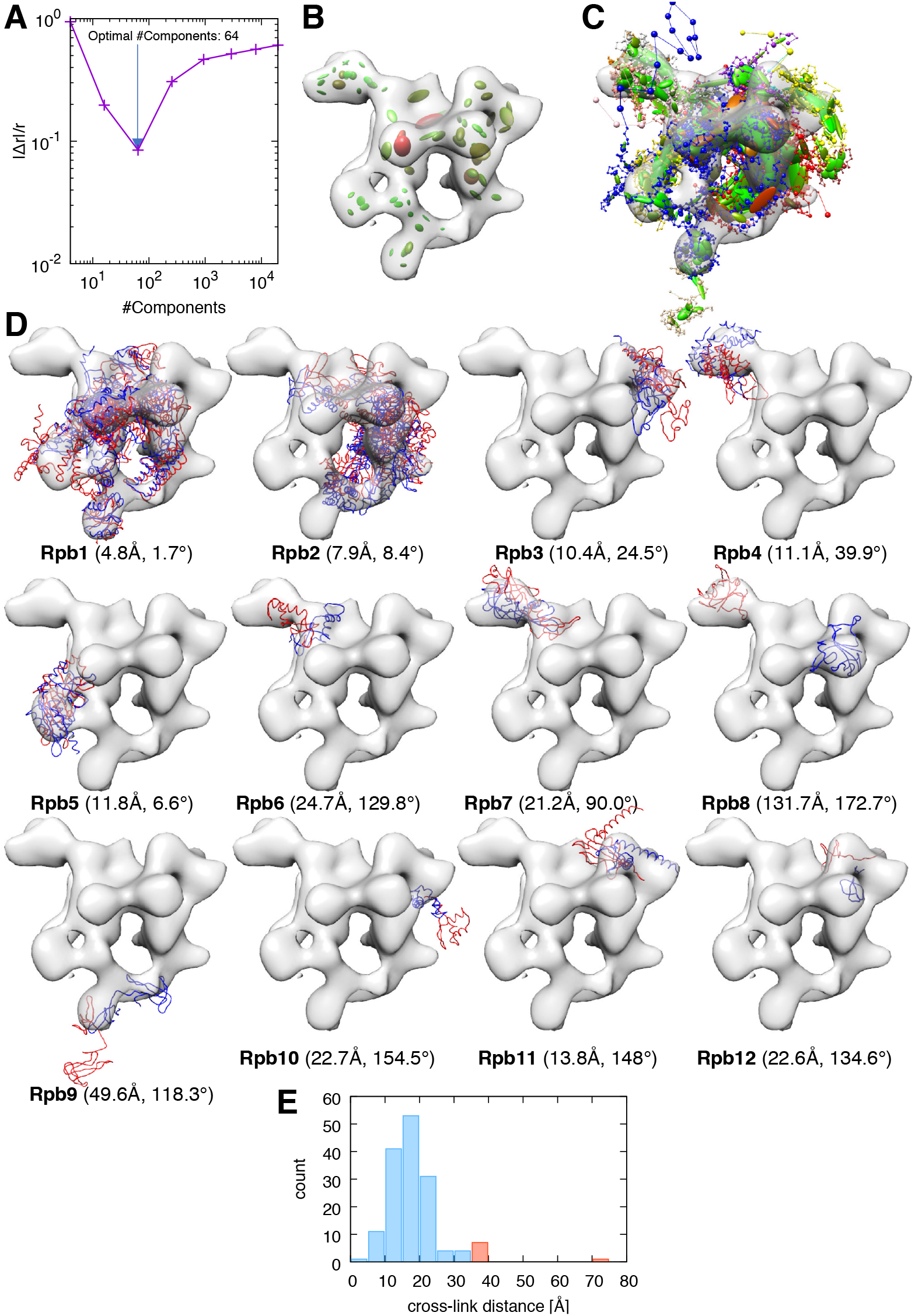
Integrative modeling of the RNA polymerase II. (A) Absolute relative deviation between data-GMM and experimental map resolutions 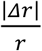 plotted as a function of the number of components of the data-GMM. The minimum (blue arrow) corresponds to the optimal number of components used in the modeling (64 Gaussians). (B) The experimental cryo-EM density map (transparent grey surface) is represented with the optimal data-GMM (colored ellipsoids). The color gradient (from green to red) is proportional to the weight *ω_D,i_* of the corresponding Gaussian. The length of the three axes and their orientation represent the 3-dimensional covariance matrix *Σ_D,i_*. (C) Representation of the best-scoring model. Coarse-grained subunits are represented by the strings of beads: the small beads and large beads represent 1- or 20-residue fragments, respectively. As for the data-GMM in panel B, the model-GMM is represented by ellipsoids. (D) All subunits of the model (red) and reference structure (PDB code 1WCM, blue) are represented along with the experimental cryo-EM map. For each panel, the name of the subunit is indicated in bold, together with the placement score of that subunit. (E) Histogram of the distance between cross-linked residues. The histogram bins corresponding to satisfied and violated cross-links are represented in blue and red, respectively.

997 of the 1000 best scoring models grouped in the first cluster. The rmsd of the best scoring model with respect to the reference structure was 32.9 Å, with a p(20) of 0.80 and a data-model CC of 0.52. The major contribution for the inaccuracy was the misplacement of subunit Rpb8. The reason for the misplacement was that Rpb8 was not cross-linked with the rest of the complex. Excluding Rpb8 from the rmsd calculation yielded a rmsd of 21.2 Å.

We analysed the position of each subunit of the complex (**Fig.6D**). Subunits 1 to 5 (81% of the mass of the complex) had a rmsd with respect to the reference structure under 20 Å. The following subunits had a rmsd over 20 Å with respect to the reference structure: Rpb6 (32.0 Å), Rpb 7 (28.1 Å), Rpb 8 (133.3 Å), Rpb9 (51.9 Å), Rpb 10 (27.3 Å), Rpb 11 (25.5 Å) and Rpb 12 (29.0 Å). Note that Rpb1 was correctly localized in the cryo-EM map but the domain corresponding to residues 1275-1733 was misplaced. Another reason for the inaccuracy is that subunits Rpb8, Rpb9, and Rpb12 were weakly cross-linked with the rest of the complex, forming 0, 1 and 3 cross-links respectively.

There was a total of 9 violated cross-links (3.5% of the total dataset), which involved the following subunits: Rpb1-Rpb1 (2 cross-links) Rpb1-Rpb2 (2 cross-links), Rpb1-Rpb4 (1 cross-link), Rpb1-Rpb6 (1 cross-link), and Rpb2-Rpb2 (3 cross-links) (**Fig.6E**).

### Integrative modeling of the exosome complex

The 10-subunit yeast exosome complex is a macromolecular machine responsible for processing and degrading RNA in eukaryotic cells (Houseley et al., 2006). To model this complex, we used the structures of all subunits of the complex in one state, the crystal structure of the RNA-bound exosome (PDB code 4IFD). We also incorporated independent data collected in another state, the low-resolution cryo-EM map of the RNA-free exosome (EMDB code 3367) (Liu et al., 2016) and a dataset of 98 cross-linked residue pairs obtained by XL-MS (Shi et al., 2015) (**Fig.7, Fig.S7, and Fig.S8**).

**Figure 7.**
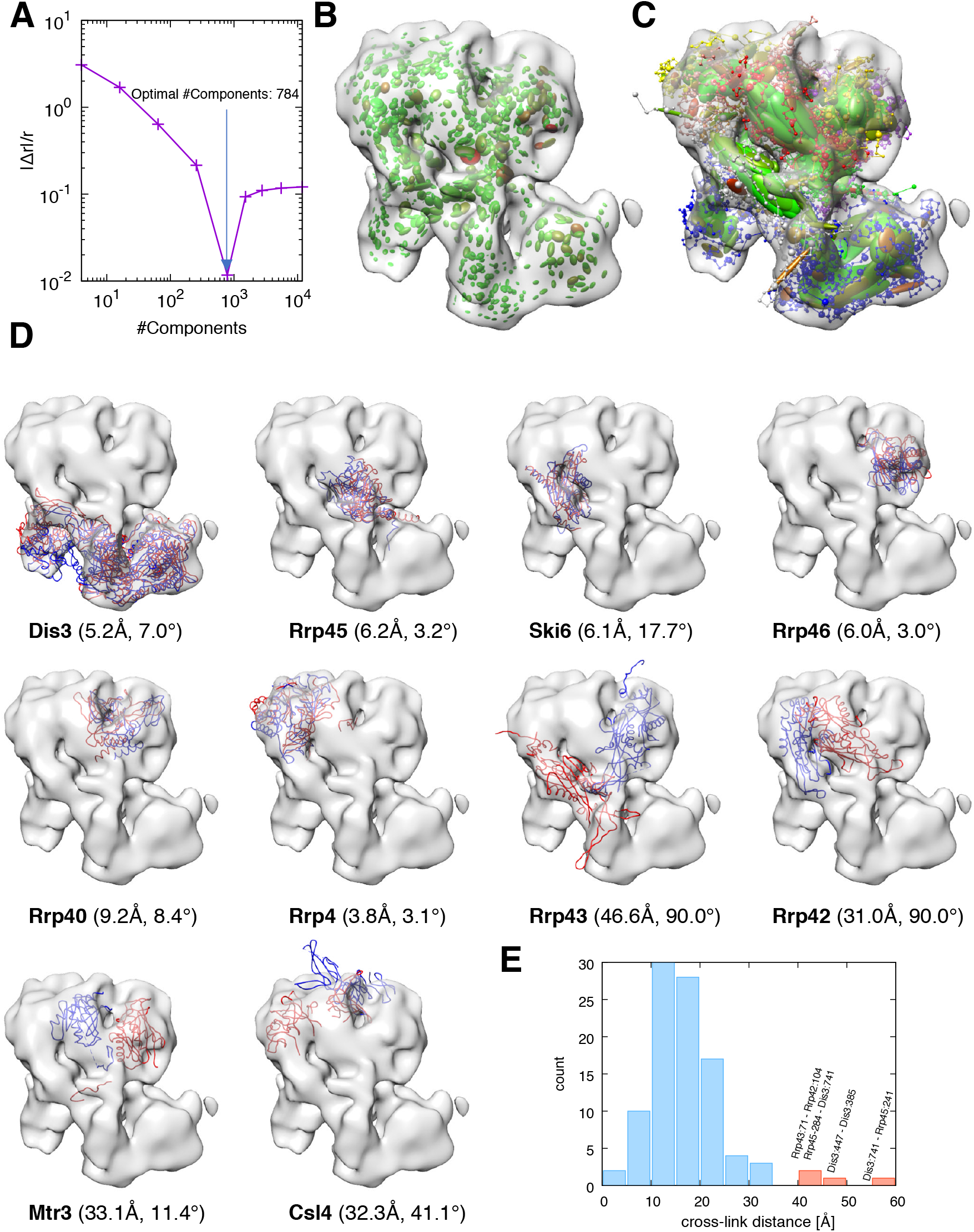
Integrative modeling of the exosome complex. We report the same information as in Fig.6 for the case of the yeast exosome complex, with the following differences: (A) the optimal number of components used in the modeling is 784; (D) the reference structure is taken from PDB code 5G06.

To the best of our knowledge, no high-resolution structure of the RNA-free 10-subunit exosome complex alone is available. We thus used the structure of the RNA-free exosome in complex with Ski7 (PDB code 5G06) as a reference to test the accuracy of our models. We expected our models to differ from the reference, especially in the region at the top where the complex interacts with Ski7. Here, for instance, when one rigidly fits the entire reference structure to the RNA-free density map, the Csl4 subunit extends outside the density map (**Fig.7D**), most likely as a consequence of the interaction with Ski7. On the other hand, the lower region in which Dis3 is located is expected to be structurally similar to the reference. In addition, cross-links were extracted from whole-cell lysate and therefore might reflect a mixture of different compositional and conformational states (Shi et al., 2015).

The 1000 best scoring models grouped into a single cluster. The rmsd of the best scoring model with respect to the reference structure was 29.4 Å, with a p(10) of 0.47 and a data-model CC of 0.79. We analysed the position of each subunit of the complex (**Fig.7D**). Strikingly, each domain of Dis3 was properly placed in its respective density region (**Fig.7D**). Subunit Csl4 was localized entirely inside the density map, at variance with the reference structure. Subunits Rrp45, Ski6, Rrp46, Rrp40, and Rrp4 occupied the correct regions of the density map. Subunits Rrp42, Rrp43, and Mtr3 were misplaced, but still occupied the upper region of the density map. The majority of the cross-links were satisfied, with measured distance between cross-linked residue pairs always below 35 Å, with a few exceptions. The cross-link between residue 71 of Rrp43 and residue 104 of Rrp42 was violated (**Fig.7E**). Three other cross-links involving Dis3 were found to be inconsistent with the cryo-EM map and therefore were violated. These distance restraints might be satisfied in the RNA-bound exosome form, which we expect to be present under the conditions in which the XL-MS data were collected.

## Discussion

A major problem in integrative structure modeling, in which data of different types are combined to model the structure of a biological complex, is to determine the relative weight of each piece of information. Inaccurate weighing results in models biased towards a particular source of data, thus reducing the accuracy of the model and under- or over-estimating its precision. To optimally weigh each piece of information, two main factors need to be considered: the accuracy or level of noise in the data and the correlation between data points.

Our Bayesian approach addresses these challenges by introducing several technical features. First, building on the *gmconvert* utility (Kawabata, 2008), we developed a divide-and-conquer strategy to efficiently compute GMM with a large number of components in order to reduce the correlation between voxels. Second, our approach accounts for the presence of variable level of noise across the experimental map and weighs each component of the GMM accordingly. Third, we created a multiscale modeling approach, as the scoring function can be adapted to any coarse-grained representation of the model and any resolution of the experimental density map. Fourth, we used a combination of flexible and rigid degrees of freedom in the modeling: each domain with a known structure is constrained into a rigid-body, while all missing parts (loops, termini, or unknown regions) are represented by flexible strings of beads. Finally, we developed an enhanced-sampling technique based on an iterative replica exchange strategy and a MC mover that randomly swaps rigid-bodies with similar shape.

### Comparison with existing approaches

To compare our approach with state-of-the-art methods for modeling macromolecular complexes using low- and intermediate-resolution cryo-EM maps, we first examined the results of the benchmark carried out with γ-TEMPy. (Pandurangan et al., 2015). This method scores the models using mutual information between model and experimental densities and uses a genetic algorithm to accelerate sampling. In the following, we compare our approach with γ-TEMPy in terms of accuracy of the scoring function, sampling efficiency, and computational performances.

In order to compare the accuracy of the two approaches, we examined our best scoring model (**Table 2**) and the γ-TEMPy high scoring model (HS in Table 1 of Ref. (Pandurangan et al., 2015)) on a subset of 9 test cases of our benchmark that were also included in the γ-TEMPy benchmark (PDB codes 2DQJ, 2BO9, 2BBK, 2GC7, 1VCB, 1TYQ, 1MDA, 1GPQ, and 1CS4). In all cases, our approach produced models that were significantly more accurate. Particularly striking is the 3-subunits complex 1VCB, which our method and γ-TEMPy modeled with rmsd of 2.2 Å and 25.3 Å, respectively.

To assess sampling efficiency, we compared our best rmsd model with the γ-TEMPy best prediction (BP in Table 1 of Ref. (Pandurangan et al., 2015)). In 8 out of 9 cases, our approach was capable to sample more native-like models. Only in the case of 1TYQ, our best rmsd model was less accurate (19.1 Å) than the BP model generated with γ-TEMPy (16.9 Å).

To assess the performances of the two approaches, we monitored the computational cost to run the two benchmarks, defined by the total number of core hours required to complete one test case. Our benchmark was executed in parallel on 48 cores on a compute cluster equipped with 2.50 GHz Intel(R) Xeon(R) E5-2670 v2 processors. The minimum, maximum, and average computational cost across all test cases was 320, 6840, and 2166 core hours, respectively. Furthermore, this computational cost depends on the number of components of the data-GMM, which is particularly advantageous with over-sampled low-resolution maps. Each computation of the γ-TEMPy benchmark was instead run on 160 cores distributed on 40 AMD 4-core 2.6 GHz processors. As the time measurements for the benchmark with 10 Å-resolution maps were not reported, we used the timings of the 20 Å-resolution benchmark to estimate the computational cost. The reported minimum, maximum, and average cost across all test cases were 640, 7840, and 2720 core hours, respectively.

We then compared our approach with other integrative modeling tools in the case of the GroEL/ES complex (**Table S3**). The accuracy of our approach was: *i)* similar to that of Attract-EM (de Vries and Zacharias, 2012), IQP (Zhang et al., 2010), and γ-TEMPy (Pandurangan et al., 2015), *ii)* superior to MultiFit (Lasker et al., 2009) and gmfit (Kawabata, 2008), and *iii)* worse than ISD (Habeck, 2017). Non-integrative modeling tools, which fit proteins into the map sequentially such as gEMfitter (Hoang et al., 2013) and PowerFit (van Zundert and Bonvin, 2015), performed equivalently or better, thanks to their exhaustive search and/or prior map segmentation. It has to be noted, however, that exhaustive search might not be amenable for large multi-component complexes, sequential fitting might bring bias to the final models, and segmentation might be incorrect.

Our approach shares the same philosophy of the Bayesian cryo-EM restraint recently developed in ISD (Habeck, 2017). However, it is distinct from it because the weight of the restraint in the ISD case is dependent on the sampling of the density map as it does not consider spatial correlation between voxels. In contrast, in our method, the number of Gaussians, and thus the weight, is independent from the grid-sampling of the density map.

### Current limitations

In the few cases in which our approach produced results of accuracy lower than the average, we identified two sources of error: a) positional ambiguity, where multiple placements result in the same score, and b) inefficient sampling of rigid body configurations in crowded environments. For example, helical bundles are difficult to model at low resolution because they only define a cylindrical shape in which two or more helices can be positioned in multiple ways. Similarly, pseudo-spherical subunits can be rotated around their center of mass or swapped with only minimal penalty. In addition, the placement of DNA helices is degenerate, because their expected density is symmetric by rotation. Finally, macromolecular complexes present a crowded environment in which sampling of rigid body configurations might be inefficient due to steric hindrance.

In applications with actual experimental density maps, we foresee an additional source of error that could affect the accuracy of the modeled complex. This error is associated with the fact that not all components of the modeled complex might have a corresponding experimental density or that the experimental density might represent more components than those explicitly modeled. Our approach currently assumes that all data-GMM components can be explained by a corresponding density of the model. This assumption is encoded in the scoring function by assigning the same total electron density to the data-GMM and the model-GMM, i.e. The two GMMs are normalized to the same value. It should be notated that this challenge, along with the two previously described, is not specific to the modeling protocol presented here, but it is faced by all the techniques to model architectures from low-resolution cryo-EM data.

### Dissemination

We implemented our modeling protocol in a series of scripts based on IMP.pmi (Webb et al., 2018), a module of the Integrative Modeling Platform (IMP, http://integrativemodeling.org) (Russel et al., 2012) that can be used to build the system representation, setup the scoring function, define the degrees of freedom to sample, and finally analyze the solutions. Our approach was also implemented in the PLUMED-ISDB module (Bonomi and Camilloni, 2017) of the open-source PLUMED library (www.plumed.org) (Tribello et al., 2014). Thanks to the differentiability of the scoring function, this implementation can be used for real-space, flexible refinement of individual models using molecular dynamics at atomistic resolution or, in combination with metainference (Bonomi et al., 2016), to model ensemble of structures representing the conformational heterogeneity hidden in low-resolution areas of atomistic density maps (Bonomi et al., 2018; Vahidi et al., 2018).

## Acknowledgements

We thank Dr. T. Huynh for assistance with the computer cluster at Institut Pasteur, and Dr. D. Saltzberg for useful discussions.

Authors contributions

S.H., M.B, R.P.performed research and analyzed the data.M.B., R.P.developed methodology.M.N.provided computing resources. All authors contributed to writing the manuscript.

## Star Methods

### Key resources table

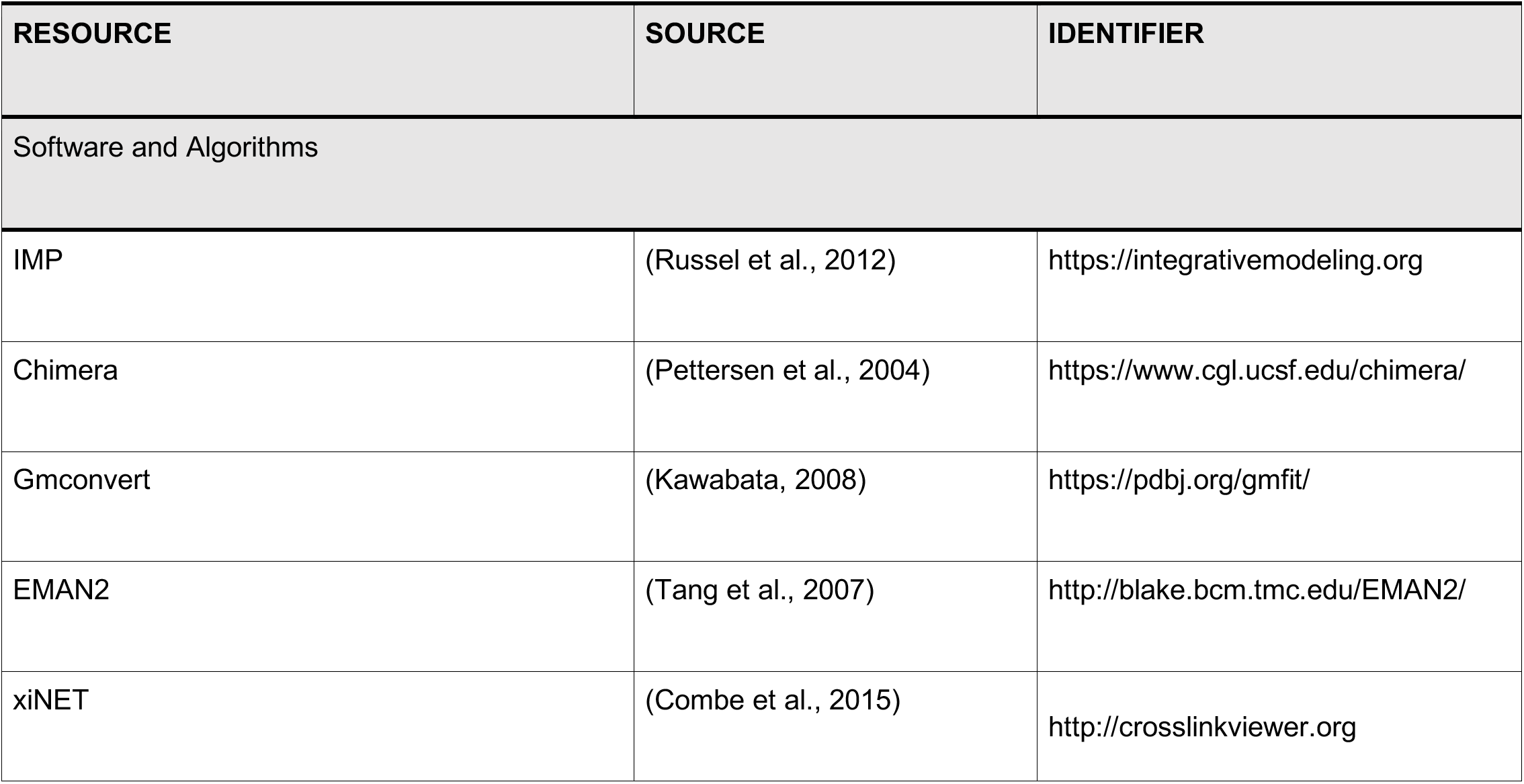

### Theory

In general terms, the Bayesian approach (Rieping et al., 2005) estimates the probability of a model, given information available about the system, including both prior knowledge and newly acquired experimental data. The posterior probability *p*(*M|D*) of model *M*, which is defined in terms of its structure *X* and other Bayesian parameters, given data *D* and prior knowledge is:

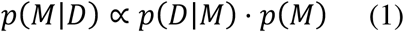

where the *likelihood function p(D|M)* is the probability of observing data *D* given *M* and the *prior p(M)* is the probability of model *M* given prior information. To define the likelihood function, one needs a *forward model f(X)* that predicts the data point that would be observed for structure *X* in the absence of experimental noise, and a *noise model* that specifies the distribution of the deviation between the experimentally observed and predicted data points. The *Bayesian scoring function* is defined as 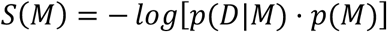 which ranks the models in the same order as the posterior probability *p(M|D)*. The prior *p(M)* includes the sequence connectivity, the excluded volume, and rigid body constraints. To compute these priors, the domains of the proteins are coarse-grained using beads of varying size. The sequence connectivity term is a sum of upper harmonic distance restraints that apply to all pairs of consecutive beads in the sequence, implied by the covalent structure of the polypeptide/polynucleotide main-chain. The excluded volume is computed from a soft-sphere potential where the radius of a bead is estimated from the sum of the masses of its residues. The structures derived from X-ray data or homology models are coarse-grained using two categories of resolution, where beads are represented either individual residues or segments of up to 10 residues. Beads can be constrained into a rigid body, in which relative distances are fixed during sampling.Alternatively, strings of beads representing parts without structural information can be flexible with respect to each other. In the following, we define the components of the Bayesian scoring function specifically for a cryo-EM density map.

*Experimental cryo-EM density map*. We represent the experimental density map *Ψ_D_* in terms of a Gaussian mixture model (GMM)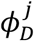 with *j* components (ie, data-GMM):

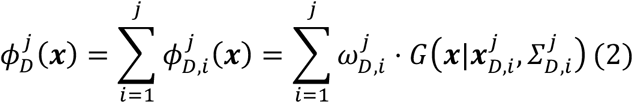

where 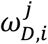 is the (normalized) weight of the *i*-th component of the GMM and *G* is a normalized Gaussian function with mean 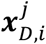 and covariance matrix 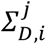

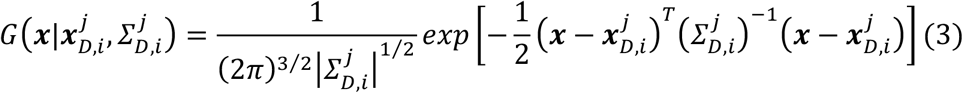

This description presents three advantages.First, it circumvents the problem of dealing with correlations in the data and noise that are typical of voxel-based representations, as each 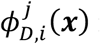 might be regarded as an independent component of the density map.Second, it provides a computationally-convenient representation of the data in terms of analytical functions.Finally, it allows representing the density map at multiple resolutions, which is exploited here to accelerate sampling of structural models compatible with the data (ie, see Model Sampling paragraph below).

The posterior probability of model *M* given the cryo-EM density map *Ψ_D_* can be written in terms of all possible GMMs that can be used to represent the data:

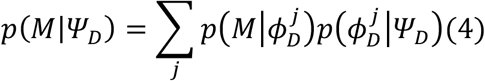

In the following, we assume that the conditional probability 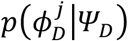 selects a single GMM *φ_D_* with *N_d_* components, which optimally represents the data. In this situation:

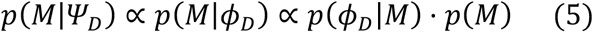

*Divide-and-conquer fit of the data-GMM*. To fit the experimental density map *Ψ_D_* with a GMM *φ_D_*, we used the Expectation Maximization algorithm implemented in the *gmconvert* software (Kawabata, 2008). This approach determines the parameters of the GMM (mean, weight, and covariance matrix of each Gaussian component) by maximizing the likelihood that the GMM density function generates the density of the voxels in *Ψ_D_*. As the resolution of the map increases, the number of Gaussians required for the GMM to accurately reproduce all the features of the experimental map increases exponentially along with the computational time and memory required to perform the fit. To overcome these challenges, we developed a divide-and-conquer approach (**Fig.2**).First, the map *Ψ_D_* is masked and all voxels with a density lower than the threshold recommended in the EMDB database are removed.Second, a recursive procedure starts from a first iteration in which the map *Ψ_D_* (**Fig.2A**) is fit with a GMM consisting of a small number of Gaussians *N_D_* (typically 2 or 4) (**Fig.2B**). Each of the components *φ_D i_* of this initial GMM is used to partition the original map into sub-maps *Ψ_D_,_i_* (**Fig.2C**):

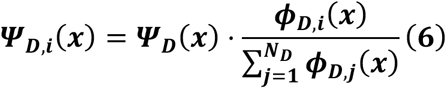

This partitioning has two properties: *a)* each sub-map selects the part of the original map that overlaps with the component (*φ_D_,_i_*); *b)* the sum of all sub-maps results in the original density map: 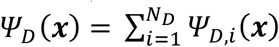 The process is repeated, and each sub-map *Ψ_D i_* is again fitted using a GMM with a small number of Gaussians *N_D_* (**Fig.2D**), dividing the process into as many branches as the number of Gaussians *N_D_*.

At each iteration the portion of the original map that is fit by a given GMM is reduced, so that a small number of Gaussians will eventually be sufficient to accurately reproduce high-resolution details.Furthermore, because of property *b)*, the *global* GMM defined by the sum of all the GMMs obtained at any given iteration also fits the original map (**Fig.2E**). This procedure is repeated until the global GMM reaches the desired accuracy (**Fig.2F**). The accuracy of the fit was defined as the correlation coefficient CC (Frenkel and Smit, 2002) between the cryo-EM density map *Ψ_D_* and the map generated by rasterizing the data-GMM into a 3D grid with the same mesh properties as the original density map (*ie*, voxel size, offsets, and box lengths) (**Fig.3A**). The CC was computed using only those voxels whose density exceeds the recommended threshold value reported in the EMDB.A given branch was stopped when the local CC between a sub-map and its GMM was greater than 0.95. Since the resolution of the original map can vary locally, individual branches will be terminated at different iterations.

This procedure generates at each step a global data-GMM with increasing number of components, thus with increasing resolution. To quantify the resolution of each of these global data-GMMs, we computed their Fourier Shell Correlations (FSCs) with respect to the original map *Ψ*_D_. By analogy with the method of the two half-maps (Rosenthal and Henderson, 2003), the resolution was defined as the inverse of the frequency at which the FSC crossed the 0.5 threshold (**Fig.S1A**).Finally, we defined the *optimal* data-GMM as the fit with resolution closest to the original map *Ψ_D_* (**Fig.S1B**). The entire process was parallelized to run efficiently on a computer cluster.

*The forward model*. We developed a forward model to compute a cryo-EM density map from a single structural model. As for the data representation above, the forward model *φ_M_* is a GMM with *N_M_* components (*ie*, model-GMM):

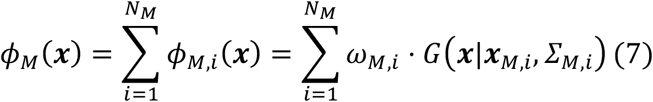

For high-resolution maps, each atom can be represented by a single Gaussian, whose parameters can be obtained by fitting the electron atomic scattering factors for a given atom type (Prince, 2004). For low-resolution maps or for an efficient initial sampling of high-resolution maps, we use a single Gaussian to represent each coarse-grained bead, with the Gaussian width proportional to the size of the bead. If multiple coarse-grained beads of the model are part of the same rigid body, the parameters of the model-GMM associated to these beads are computed by applying the Expectation-Maximization algorithm to the positions of the centers of the beads, weighed by their mass.

*The noise model*. At variance with our previous effort in modeling cryo-EM data (Robinson et al., 2015), in this approach we will not use the global correlation coefficient (CC) as measure of agreement between predicted and observed density maps, but a likelihood obtained from the product of functions of local cross-correlation-like terms, as explained below.First, we define the global overlap between model and data density maps as:

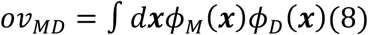

The standard CC can then be expressed in terms of the overlap functions as (Robinson et al., 2015):

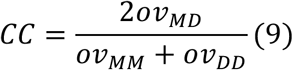

Notably, maximum correlation is obtained with maximum overlap *ov*_MD_, since the quantities at the denominator of Eq.9 do not depend on the coordinates of the particles in the structural model.

The global overlap *ov_MD_* can be expressed in terms of local overlaps *ov_MD k_* between model and the *k-* th component of the data-GMM *φ*_D_,_k_:

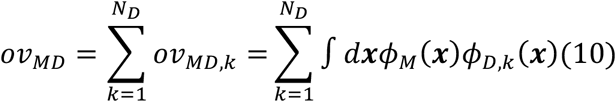

Each local overlap measures the agreement of the model with the part of the experimental density map represented by a component of the data-GMM. Because *φ_M_* is also a GMM, we can write the local overlap as the sum of overlaps for the individual components:

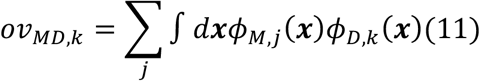

where the overlap between two Gaussians *φ_M_,_j_* and *φ_D_,_k_* is given by:

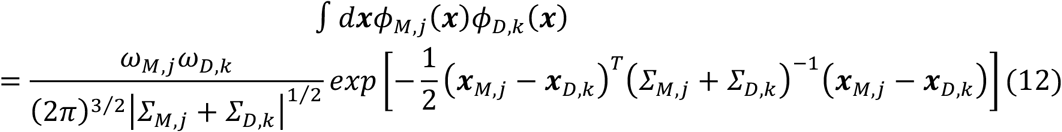

We treat the *N_D_* individual components of the data-GMM as independent pieces of information and express the data likelihood in terms of local overlaps *ov*_MDk_, using a log-normal noise model:

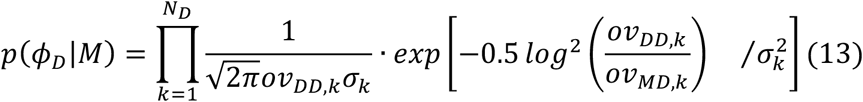

where *σ_k_* is the unknown tolerance associated with the *k-th* component of the data-GMM and *ov_DD__k_* is the overlap of the *k*-th component with the entire data-GMM. It should be noted that a GMM represents the experimental data with less correlated components compared to the voxel representation.However, we expect a residual correlation amo
ng GMM components, which we explicitly neglect when writing Eq.13.

*Marginal likelihood*. For simplicity, in the following we assume that different parts of the map have the same tolerance *σ* and we marginalize this variable using an uninformative Jeffreys prior 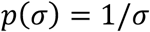 The resulting marginal data likelihood can be written as:

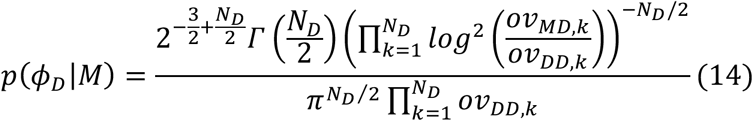

Alternatively, one can assume a variable level of noise in the map and marginalize each *σ_k_* using a Jeffreys prior. The marginal likelihood in Eq.14 is maximized when the local overlap *ov_MD_,_k_* reproduces the overlap *ov_DD_,_k_* for all data-GMM components (**Fig.S2**).

*Bayesian scoring function*. Omitting constant quantities, the final Bayesian scoring function for a fit of a model to a cryo-EM map can be written as:

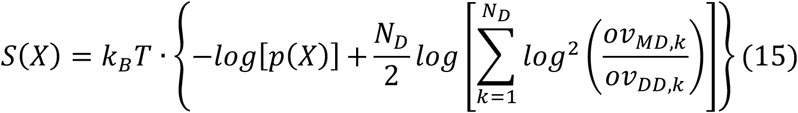

where *p(X)* is the structural prior and depends on the resolution of the model. In our coarse-grained representation, p(X)is the sum of an excluded volume potential to avoid steric clashes and a sequence connectivity restraint.

Importantly, the number of components *N_D_* of the data-GMM determines the overall weight of the cryo-EM restraint by increasing the number of log-harmonic functions. On the other hand, the weight is less sensitive to the number of components of the model-GMM. As a consequence, the data-GMM has to be rigorously fit to the experimental density map with the divide-and-conquer approach, while there are no strict guidelines for the maximum number of components of the model-GMM.Here, we followed a parsimonious approach and we empirically chose the number of components in the model-GMM to match the number of Gaussians per unit of mass in the data-GMM. With high-resolution density maps and atomistic models, we expect to use one component of the model-GMM per heavy atom of the system.

### Benchmark of the divide-and-conquer fit of the data-GMM

We assessed the accuracy of our divide-and-conquer approach to computing a data-GMM by using 20 experimental density maps of protein complexes at different resolutions, ranging from 3.6 to 25 Å (**Table 1**). We used the divide-and-conquer approach described above to obtain GMMs of each map with a number of Gaussians varying from 16 to 16384. At each step of the divide-and-conquer each sub-map was fit using a GMM with 4 components.

### Benchmark of the modeling protocol

*Data generation*. We benchmarked our modeling protocol using 21 protein/DNA complexes consisting of 2 to 7 subunits (Velazquez-Muriel et al., 2012) (**Table 2 and Fig.4**). For each of these complexes, we generated a simulated cryo-EM density map, using the structures extracted from the PDB. We used one Gaussian for every 1.090 kDa of assembly mass, which corresponded approximately to the mass of 10 residues and resulted in a resolution of approximately 10 Å, as obtained by extrapolation from the stretched-exponential regression (**Fig.3B**). For example, the human transferrin receptor complex (PDB code 1SUV) (Cheng et al., 2004) consists of 6 subunits and has a molecular mass of 290 kDa. Therefore, the simulated map was determined using 262 Gaussians. The simulated GMMs were generated from the reference structures using the program *gmconvert* (Kawabata, 2008).

*Subunits representation and forward model*. Molecules (protein and DNA chains) were represented by a set of spherical beads, each with the volume of the corresponding residue. When available, the positions of beads were obtained from the PDB structures and constrained into one or more rigid bodies. Missing regions were constructed as strings of flexible coarse-grained beads. When molecules were intertwined or if a molecule was composed of structurally independent domains, we defined several rigid bodies, one for each domain.Furthermore, in some cases, two domains belonging to distinct molecules were merged into the same rigid body, such as the DNA double-strands in 3V6D and 2Y7H or the helical bundle of 3PUV. The model-GMM was computed as follows.First, for each rigid body defined above, we computed a GMM based on the corresponding atomic coordinates using the implementation of the expectation-maximization algorithm available in the *scikit-learn* python library (Pedregosa et al., 2011). The number of Gaussians of a model-GMM was determined by dividing the molecular weight of the corresponding rigid body by the average weight of a 10-residue peptide (1.09kDa). The center and covariance matrix rotation of each Gaussian were constrained into the corresponding rigid body.Second, each flexible bead was treated as an individual spherical Gaussian.

*Model sampling*. The initial positions and orientations of rigid bodies and flexible beads were randomized. The generation of structural models was performed using MC coupled with replica exchange (Swendsen and Wang, 1986).48 replicas were used to cover a temperature range between 1 and 2.5 score units (SU). Intermediate temperatures followed a geometrical progression. Each MC step consisted of: A) a series of random transformations of the positions of the flexible beads and the rigid bodies, B) rigid body transformation of the whole system, and C) rigid-body swapping moves. In (A), each individual flexible bead and rigid body was translated in a random direction by up to 4 Å, and each individual rigid body was rotated around its center of mass by up to 0.04 radians about a randomly oriented axis. In (B), a rigid-body transformation was applied to the whole system. In (C) we swapped the position and orientation of two rigid-bodies, randomly chosen among those with similar shape, to allow efficient sampling of alternative conformations equally consistent with the data. The shape similarity was assessed by computing the rmsd of the inertia moments of the two rigid-bodies. Each MC step was accepted or rejected according to the Metropolis criterion.

Our scoring function becomes more rugged at higher resolutions. In fact, as discussed above, the number of components for the data-GMM *(N*_D_) increases with the resolution, and at the same time their variance decreases to better describe high-frequency features. As a consequence, the log-square score term (Eq.15) becomes more peaked, thus increasing the frustration of the total score. To alleviate this issue, we implemented an iterative sampling procedure (**Fig.S3**). The idea is to progressively increase *N_d_* using all fits obtained at different stages of the divide-and-conquer procedure, from the minimum (*ie*, 4) to the *N_D_* of the optimal data-GMM. In each iteration, we: 1) sampled the models at a given *N_d_;* 2) generated a pool of initial models (seeds) for the next iteration; and 3) incremented *N*_D_. During step (1), we produced an ensemble of 12,000 models using the MC and replica exchange protocol described above. After extracting the 100 best scoring models of the resulting ensemble, we identified a subset of models as structurally diverse as possible, using rmsd criterion. The number of models in the subset was constrained to the number of replicas (ie, 48). In the first iteration *(N_D_* =4), the score-landscape is shallow, which allows the system to explore a large variety of conformations. As *N_D_* increases, the structural variability among seeds is reduced.

Finally, to assess sampling exhaustiveness, at the end of each iterative modeling run we analyzed the agreement of the best scoring models with the cryo-EM map by computing the cross correlation between the model-GMM and data-GMM. If the resulting cross correlations were below 0.7, we started another iteration run. The threshold of 0.7 was chosen by experience, as we noticed that lower cross correlation coefficients usually indicate poor agreement of the model with the cryo-EM data with clearly misplaced subunits. This procedure was applied to all cases for which the simple replica exchange above did not yield satisfying results (PDB codes 1MDA, 1SUV, 1TYQ, and 3PUV in the synthetic benchmark and the application to the RNA polymerase II and exosome complexes).

*Analysis*. All models produced by the modeling protocol described above were ranked by score, and the 1000 best scoring models were considered for further analysis. The accuracy of the fit was assessed by computing a series of structural metrics, namely the rmsd, p(10), the correlation coefficient between the model- and data-GMMs, as well as the average placement score of the best scoring model. These metrics are defined in the next paragraph.

### Structural metrics

To compare two models, we used several metrics, including the rmsd of residue positions, p(10), the Average Placement Score (APS), and the data-model correlation coefficient CC. The rmsd of residue positions was defined as the rmsd between the positions of corresponding centers of the coarse-grained spheres in two structures, without structural alignment. When multiple copies of the same protein were present, the rmsd was defined as the minimum rmsd across all possible assignments of the identical components.p(10) was defined as the percentage of residues whose deviation between the two structures is lower than 10 Å. The Placement Score of the model is a two-number metrics that measures the translation and the rotation needed to optimally align each subunit of the model to a reference. The APS is average of the Placement Score calculated over all subunits and weighted by the number of residues. The data-model correlation coefficient (CC) was defined by Eq.9 and quantifies the agreement of the model with the data.

*Clustering*. For each complex, the 1000 best scoring models selected for analysis were clustered using a hierarchical clustering approach (Johnson, 1967).Initially, all 1000 models were placed in a list *L* of models not yet clustered. Then, we applied the following iterative procedure:

1. The best scoring model *m0* from the list *L* was selected to define a new cluster *Ck* and removed from *L*.
2. All models *mi* from *L* with rmsd from *m0* lower than 10 Å were defined as members of the cluster *Ck* and removed from *L*.
3. We iterated step 1 and 2 until all models were clustered into spheres of radius equal to 10 Å.

At the end of this iterative procedure, we merged all those pairs of clusters that contained at least two elements within 10 Å one from each other. By construction, the first cluster produced by the algorithm (labelled as C1) contained the best scoring model.

### Modeling of the GroEL/ES complex

We modeled the architecture of the 21-subunit GroEL/ES ADP-bound complex using cryo-EM data at 23.5 Å resolution (EMDB code 1046) (**Fig.5**). The GroEL/ES complex consists of 2 sequences, the chaperonin GroEL and the cochaperonin GroES, with a stoichiometry 14:7. The 14 copies of GroEL have two distinct structures, named GroEL-cis and GroEL-trans. The 7 GroES, 7 GroEL-cis, and 7 GroES-trans are arranged in a C7 symmetry, each one occupying one centro-symmetric ring (**Fig.5**). The optimal data-GMM contained 256 components (**Fig.S4A**). We followed the modeling protocol and the model representation used for the benchmark with synthetic cryo-EM maps (**Fig.S4B**). The coordinates of the beads used to represent our system were obtained from PDB code 1AON (Armache et al., 2005). Each protein was constrained into a rigid-body based on the crystallographic structure, and a C7 symmetry constraint is applied. The number of residues per coarse-grained bead was set to 20, and the number of residues per component of the model-GMM was set to 10. The quality of the resulting models was assessed using the same structural metrics as in the benchmark with synthetic data, using the structure of PDB code 1AON as reference.

### Integrative modeling of the RNA polymerase II complex

We modeled the architecture of the 12-subunit RNA polymerase II using cryo-EM data at 20.9 Å resolution (EMDB code 1883) (Czeko et al., 2011) and two datasets of 108 (Chen et al., 2010) and 157 (Robinson et al., 2015) cross-links (107 inter- and 158 intra-molecular) (**Fig.6 and Fig.S5**). The RNA polymerase II complex consists of 12 subunits, named Rpb1 to Rpb12. The optimal data-GMM contained 64 components (**Fig.6A**). We followed the modeling protocol and the model representation used for the benchmark with synthetic cryo-EM maps (**Fig.S6**). The coordinates of the beads used to represent our system were obtained from PDB code 1WCM (Armache et al., 2005). Based on a prior domain analysis of Rpb1 and Rpb2, we constrained the coordinates of these two large subunits into four rigid-bodies corresponding to: 1) residues 1141-1274 of Rpb1, 2) residues 1275-1733 of Rpb1, 3) residues 1-1102 of Rpb2, and 4) residues 1-1140 of Rpb1 together with residues 1103-1224 of Rpb2. The number of residues per coarse-grained bead was set to 20, and the number of residues per component of the model-GMM was set to 10. The quality of the resulting models was assessed using the same structural metrics as in the benchmark with synthetic data, using the structure of PDB code 1WCM as reference. The XL-MS data was encoded using a previously developed Bayesian scoring function (Shi et al., 2015).

### Integrative modeling of the exosome complex

We modeled the architecture of the 10-subunit yeast exosome complex using cryo-EM data at 11.5 Å resolution (EMDB code 3367) (Liu et al., 2016) and a dataset of 98 cross-links (26 inter- and 72 intramolecular) (Shi et al., 2015) (**Fig.7 and Fig.S7**). The exosome complex consists of a core complex of 9 proteins (Csl4, Mtr3, Rrp4, Rrp40, Rrp42, Rrp43, Rrp45, Rrp46, and Ski6), and an RNase protein (Dis3). The top of the core complex recruits RNAs that are then transferred to Dis3 through a central channel in the core complex. The optimal data-GMM contained 784 components (**Fig.7A**). We followed the modeling protocol and the model representation (**Fig.S8**) used for the benchmark with synthetic cryo-EM maps, with few variations. We split the largest subunit, Dis3, into three rigid-bodies, corresponding to residues 1-237, 238-471, and 472-1001, which is the domain organization of this subunit based on its structure in PDB code 4IFD (**Fig.S7**) (Makino et al., 2013). The quality of the resulting models was assessed using the same structural metrics as in the benchmark with synthetic data. The only difference was that the reference structure used to compute the accuracy (PDB code 5G06) was different from the structure used to initialize the positions of beads in rigid-bodies (PDB code 4IFD) (Liu et al., 2016). As for the modeling of RNA polymerase II, XL-MS data was encoded using a Bayesian scoring function (Shi et al., 2015).

### Quantification and statistical analysis

The analysis of the results was performed using the IMP.pmi module of the IMP software (Russel et al., 2012).

### Data and software availability

Our Bayesian approach for cryo-EM data is implemented in the *Integrative Modeling Platform* (IMP) (Russel et al., 2012), which is freely available at https://integrativemodeling.org. In particular, the representations and degrees of freedom of each complex were encoded in a standard way using the IMP.pmi topology tables.

